# NSE5 subunit interacts with distant regions of the SMC arms in the *Physcomitrium patens* SMC5/6 complex

**DOI:** 10.1101/2024.01.12.574989

**Authors:** Jitka Vaculíková, Marcela Holá, Barbora Králová, Edit Lelkes, Barbora Štefanovie, Radka Vágnerová, Karel J. Angelis, Jan J. Paleček

**Affiliations:** National Center for Biomolecular Research, Faculty of Science, Masaryk University, Kamenice 5, 62500 Brno, Czech Republic; Institute of Experimental Botany, Czech Acad Sci, Na Karlovce 1, 16000 Prague, Czech Republic; Mendel Centre for Plant Genomics and Proteomics, Central European Institute of Technology, Masaryk University, Kamenice 5, 62500 Brno, Czech Republic

**Author notes:** Corresponding addresses, National Center for Biomolecular Research, Faculty of Science, Masaryk University, Kamenice 5, 62500 Brno, Czech Republic, Institute of Experimental Botany of the Czech Academy of Sciences, Na Karlovce 1, CZ-16000, Prague, Czech Republic.

**Keywords:** SMC5/6 complex, NSE5/SNI1/SLF1/SIMC1, NSE6/ASAP1/SLF2/KRE29, DNA damage repair, rDNA stability, moss plant development

## Abstract

Structural Maintenance of Chromosome (SMC) complexes play roles in cohesion, condensation, replication, transcription, and DNA repair. Their cores are composed of SMC proteins with a unique structure consisting of an ATPase head, long arm, and hinge. SMC complexes form long rod-like structures, which can change to ring-like and elbow-bent conformations upon binding ATP, DNA and other regulatory factors. These SMC dynamic conformational changes are involved in their loading, translocation, and DNA loop extrusion. Here, we examined the binding and role of the PpNSE5 regulatory factor of *Physcomitrium patens* PpSMC5/6 complex. We found that the PpNSE5 C-terminal half (aa230-505) is required for binding to its PpNSE6 partner, while the N-terminal half (aa1-230) binds PpSMC subunits. Specifically, the first 71 amino acids of PpNSE5 were required for binding to PpSMC6. Interestingly, the PpNSE5 binding required the PpSMC6 head-proximal joint region and PpSMC5 hinge-proximal arm, suggesting a long distance between binding sites on PpSMC5 and PpSMC6 arms. Given the long distance between these PpSMC sites and the size of PpNSE5, we hypothesize that PpNSE5 either links two antiparallel SMC5/6 complexes or binds one SMC5/6 in elbow-bent conformation.

In addition, we generated the *P. patens* mutant lines (*Ppnse5KO1* and *Ppnse5KO2*) with CRISPR/Cas9-integrated stop codons in PpNSE5. The *Ppnse5KO1* mutant line with an N-terminally truncated version of PpNSE5 (starting from an alternative aaMet72) exhibited DNA repair defects while keeping a normal number of rDNA repeats. As the first 71 amino acids of PpNSE5 are required for PpSMC6 binding, our results suggest the specific role of PpNSE5-PpSMC6 interaction in DNA repair. Altogether, our study suggests that PpNSE5 binding to distant regions of the PpSMC5 and PpSMC6 arms serves a specific role in loading at DNA lesions.

## INTRODUCTION

Structural Maintenance of Chromosome (SMC) complexes are molecular machines that organize chromatin and regulate its dynamics (Uhlmann, 2016, Davidson and Peters, 2021). Best characterized are the eukaryotic cohesin and condensin that organize chromosomal DNAs into loops during interphase and mitosis, respectively. Although the SMC5/6 complex can also extrude loops (Pradhan *et al.*, 2023), its function(s) is less understood. SMC5/6 was implicated in DNA repair, replication fork restart, rDNA repeat stability maintenance, segregation, transcription, and viral restriction, but the interrelations between these functions are unclear (Aragón, 2018, Palecek, 2019).

The core of the SMC complexes is formed by two SMC proteins, kleisin, and kleisin-associated subunits (Hassler *et al.*, 2018). The SMC proteins fold into a unique structure consisting of an ATPase head at one end and a long coiled-coil arm separating the head from the hinge at the other. The hinge domains dimerize SMC molecules, and arms closely align from hinge to head in a “zipped-up” conformation, resulting in a rod-like structure tens of nanometers long. Upon ATP binding to ATPase heads, the arms “unzip” and adopt ring-shaped conformation. The other core subunits, kleisin (NSE4) and kleisin-associated (KITE: NSE1 and NSE3), bind DNA and regulate the rod-to-ring SMC5-SMC6 dynamics (Palecek and Gruber, 2015, Zabrady *et al.*, 2016, Vondrova *et al.*, 2020, Yu *et al.*, 2022). These rod-to-ring arm transitions of SMC complexes are possible via discontinuities in their coiled-coil structure. The discontinuities also allow a sharp bending of arms at “elbows”, bringing hinges close to the head area (Lee *et al.*, 2020, Petela *et al.*, 2021). Such bending and the opening of the hinges are considered key steps in the DNA loading; e.g., cohesin bending positions head-proximal Scc2 loading factor close to SMC hinges (Collier *et al.*, 2020, Shi *et al.*, 2020, Higashi *et al.*, 2021, Collier and Nasmyth, 2022). Upon SMC complex loading, the ATP binding and hydrolysis drive complex translocation along DNA or loop extrusion (Ganji *et al.*, 2018, Davidson *et al.*, 2019, Pradhan *et al.*, 2023). Interestingly, SMC5/6 translocates as a monomer, while loop extrusion requires dimerization of complex.

Multiple functions of the SMC5/6 complexes are facilitated by various factors (Mahrik *et al.*, 2023), of which the NSE5-NSE6 dimer is the best characterized at the molecular level (Oravcová and Boddy, 2019). Yeast NSE5-NSE6 dimers interact with BRCT domain-containing proteins Brc1/Rtt107, which in turn bind to damage-induced phosphorylated H2A and load SMC5/6 at DNA lesions (Williams *et al.*, 2010, Li *et al.*, 2012). In humans, NSE5A/SLF1 ortholog contains the BRCT domain, which recognizes damage-induced phosphorylation of RAD18 and targets SMC5/6 to DNA lesions (Räschle *et al.*, 2015). Another human NSE5B/SIMC1 ortholog contains SUMO interacting motifs (SIMs), which bind SUMOylated proteins in viral replication centres and facilitate SMC5/6 role in viral restriction (Oravcova *et al.*, 2022). Despite very low sequence similarities between yeast and human NSE5 factors, they share some common features (particularly, they dimerize with NSE6 and directly or indirectly associate with SMC5-SMC6 subunits). The way they associate with SMC5-SMC6 proteins is somehow divergent (Palecek *et al.*, 2006, Gutierrez-Escribano *et al.*, 2020, Taschner *et al.*, 2021, Yu *et al.*, 2021). So far no direct binding of human NSE5 orthologs to SMC proteins was found (Adamus *et al.*, 2020, Oravcova *et al.*, 2022), and reports on yeast orthologs caused controversy. Crosslinking studies placed yeast scNSE5 primarily to SMC head-proximal areas (Gutierrez-Escribano *et al.*, 2020, Taschner *et al.*, 2021, Yu *et al.*, 2021), while pull-down experiments suggested its association with SMC hinges (Duan *et al.*, 2009).

To bring new insight into the NSE5 features and controversy, we analysed PpNSE5 interactions with PpNSE6 and PpSMC subunits of *Physcomitrium patens*. We found PpSMC5 hinge-proximal and PpSMC6 head-proximal regions, which are tens of nanometres apart in SMC5/6 rod-like structure, essential for PpNSE5 binding. To interpret these findings, we proposed that NSE5-NSE6 dimers either bridge these regions between two antiparallel SMC5/6 complexes or bind them when they are next to each other within the elbow-bent SMC5/6 structure. To analyse the role of PpNSE5 binding to PpSMC6, we generated the *P. patens* mutant line (KO1) carrying a truncated version of PpNSE5, missing amino acids specifically required for PpSMC6 binding. This *Ppnse5KO1* mutant line exhibited DNA repair defects with the rDNA number of repeats unaffected, suggesting the specific role of PpNSE5-PpSMC6 interaction in DNA repair.

## RESULTS

### *Physcomitrium patens* PpNSE5 region aa230-505 is essential for binding to PpNSE6

Recently, we characterized the moss PpNSE6 subunit of SMC5/6 and its binding to the PpNSE5 partner (Lelkes *et al.*, 2023). Here, we complemented this analysis with a study of the binding properties of PpNSE5. First, we cloned its N- and C-terminally truncated PpNSE5 fragments to yeast 2-hybrid (Y2H) vectors (Fig. 1A). We found Gal4BD-PpNSE5(aa190-526) and (aa230-526) fragments binding to full-length (FL) Gal4AD-PpNSE6(aa1-480), while Gal4BD-PpNSE5(aa265-526) was unable to bind (Fig. 1B, lanes 1-4). Similarly, Gal4BD-PpNSE5(aa1-505) was able to bind PpNSE6, while fragments aa1-485 and aa1-455 could not (Fig. 1B, lanes 5-7). Notably, the binding of PpNSE5(aa230-526) and (aa1-505) fragments was weaker than FL PpNSE5 (Fig. 1B, compare 0.2 mM AT middle and 5 mM AT bottom panels). These results suggest the essential role of the C-terminal half (aa230-505) and the stabilizing effect of N-(aa190-230) and C-terminal (aa505-526) flanking sequences.

**Figure 1:**
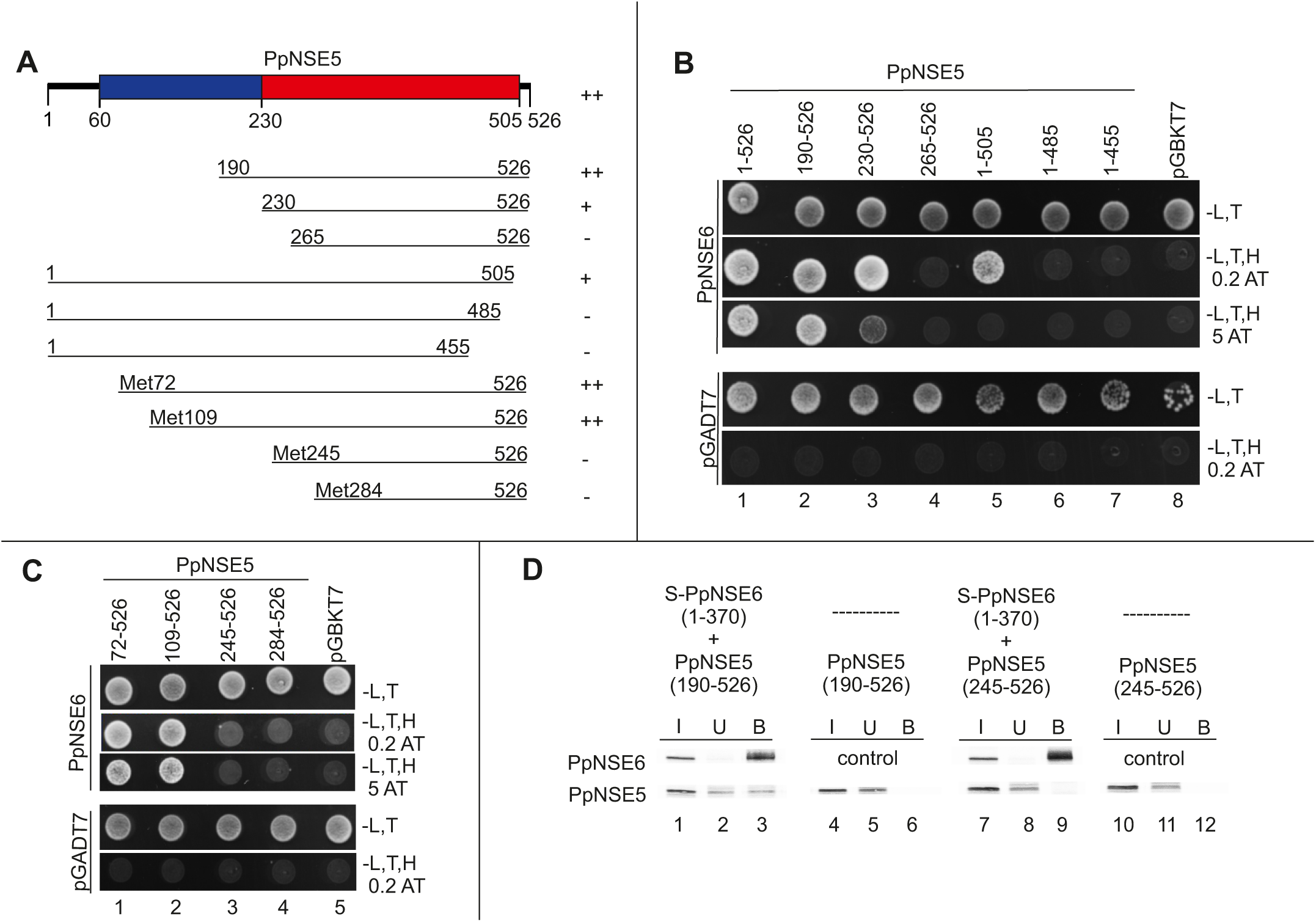
Analysis of the PpNSE5-PpNSE6 interaction. **(A)** Schematic representation of the different PpNSE5 fragments. The unstructured N-terminal part (aa1-60) is shown as a black line; two parts of the structured region are coloured in blue (aa60-230) and red (aa230-505), respectively (Fig. S1A). The PpNSE5 fragments borders were designed based on secondary and tertiary structure predictions (Fig. S1A) or the positions of native methionines. The summary of Y2H results (from panels B and C) is on the right: ++, strong interaction; +, weak interaction; –, no interaction. **(B)** The binding of N- and C-terminally truncated Gal4BD-PpNSE5 constructs to Gal4AD-PpNSE6(aa1-480) was tested in Y2H. Only FL PpNSE5 and PpNSE5(aa190-526) constructs strongly bound PpNSE6 (lanes 1 and 2, lower panel, growth on 5 mM AT plate), while binding of PpNSE5(aa230-526) and (aa1-505) was weaker (lanes 3 and 5, middle panel, growth on 0.2 mM AT plate only). PpNSE5(aa265-526), (aa1-485), and (aa1-455) fragments did not bind PpNSE6 (lanes 4, 6, and 7), suggesting the essential role of aa230-505 for PpNSE5-PpNSE6 interaction. **(C)** Fragments starting from native methionines Gal4BD-PpNSE5(aa72-526) and (aa109-526) strongly bound Gal4AD-PpNSE6(aa1-480), while Gal4BD-PpNSE5(aa245-526) and Gal4BD-PpNSE5(aa284-526) did not. All Y2H protein-protein interactions were scored by the growth of the yeast PJ69-4 transformants on the plates without Leu, Trp, His (-L,T,H), and with the indicated concentration of 3-Amino-1,2,4-triazole (AT). Control plates were lacking only Leu and Trp (-L,T). Empty pGBKT7 and pGADT7 vectors were used as negative controls. **(D)** The Y2H results were verified using *in vitro* pull-down assays. The plasmids pGBKT7-PpNSE5(aa190-526), pGBKT7-PpNSE5(aa245-526), and pET-Duet-PpNSE6(aa1-370)-StrepTwin-Stag were used for *in vitro* expression, radiolabelled proteins were mixed (as indicated), and added to the protein-S beads. In the control experiments, PpNSE5 proteins were applied to empty beads. The input (I), unbound (U), and bound (B) fractions were separated on 12% SDS-PAGE and transferred to nitrocellulose membranes. The blots were scanned for autoradiography.

As our STOP-codon CRISPR/Cas9 strategy produced an N-terminally truncated PpNSE5 fragment translated from an alternative start site (see below), we also prepared fragments starting from native methionines (Met72, Met109, Met245 and Met284; Fig. 1A). The Gal4BD-PpNSE5(aa72-526) and (aa109-526) fragments bound FL PpNSE6 with an affinity similar to FL PpNSE5, while the PpNSE5(aa245-526) and PpNSE5(aa284-526) fragments lost the binding completely (Fig. 1C, lanes 1-4). These data confirm the above conclusion that the N-terminal part of PpNSE5 is dispensable, while amino acids from position aa230 are essential for the PpNSE5-PpNSE6 interaction.

To verify our Y2H results, we expressed the PpNSE5(aa190-526) and PpNSE5(aa245-526) fragments *in vitro* and used them in a pull-down assay (Fig. 1D; (Lelkes *et al.*, 2023)). The Stag-PpNSE6(aa1-370) specifically precipitated PpNSE5(aa190-526), while not precipitating PpNSE5(aa245-526) (Fig. 1D, lanes 3 and 9), thus confirming the Y2H results and role of the C-terminal half of PpNSE5 in PpNSE6 binding.

### The first 71 amino acids of PpNSE5 are required for binding to PpSMC6

Yeast NSE5 subunits bind directly to SMC arms (Palecek *et al.*, 2006, Li *et al.*, 2023), while there is no evidence for the direct binding of human hsNSE5 orthologs (hsNSE5A/SLF1 or hsNSE5B/SIMC1) to SMC subunits (Adamus *et al.*, 2020, Oravcova *et al.*, 2022). Therefore, we tested the above constructs (Fig. 1A) and other PpNSE5 fragments (Fig. 2) against PpSMC5 and PpSMC6 constructs (Lelkes *et al.*, 2023). Surprisingly, only PpNSE5(aa1-230) fragment bound PpSMC5 and PpSMC6 (Fig. 2A). To verify this unexpected result, we used His-T7tag-PpNSE5(aa1-230) protein in the pull-down assay against PpSMC5 and PpSMC6 arm constructs (see below). His-T7tag-PpNSE5(aa1-230) was expressed in bacteria, prebound on anti-T7tag agarose beads, and then either PpSMC5 or PpSMC6 *in vitro* expressed proteins were added. PpNSE5 interacted specifically with both PpSMC constructs (Fig. 2B, lane 3), confirming direct PpNSE5 binding to PpSMC5 and PpSMC6 subunits.

**Figure 2:**
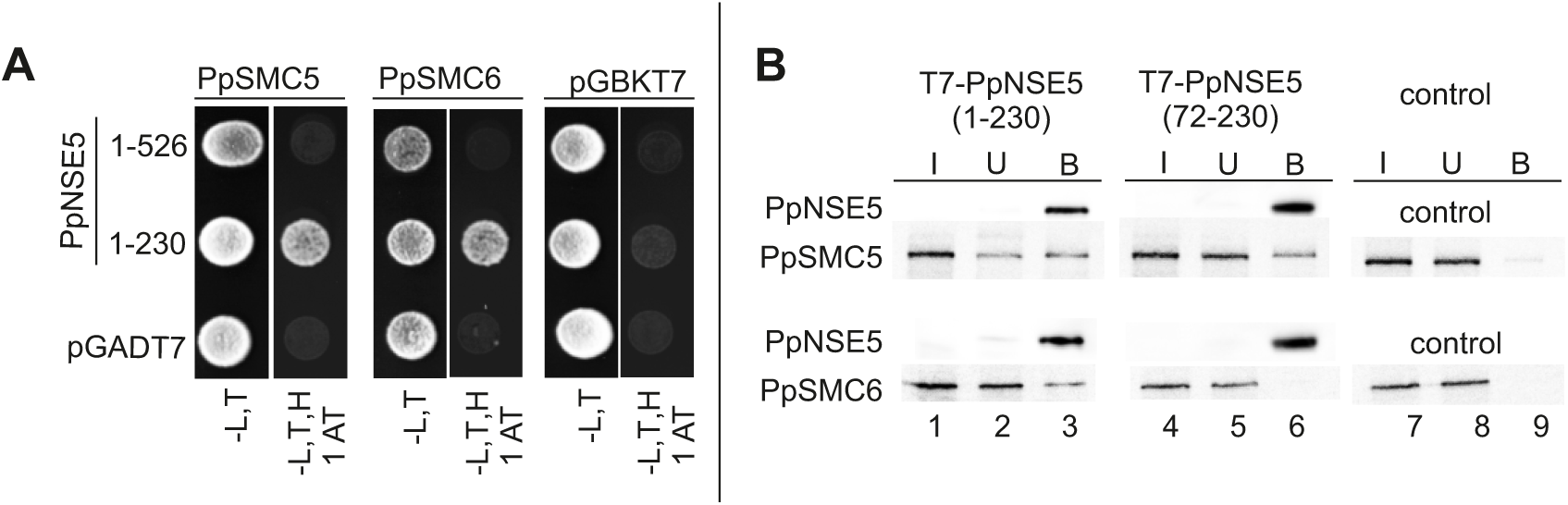
Analysis of PpNSE5 binding to SMC subunits. **(A)** The Gal4AD-PpNSE5 fragments were tested in Y2H for protein-protein interactions with Gal4BD-PpSMC5(aa201-890) (left panel) and Gal4BD-PpSMC6(aa226-955) (middle panel). Only the N-terminal PpNSE5(aa1-230) fragment bound PpSMC5 and PpSMC6. **(B)** The PpNSE5 interactions were further analysed using *in vitro* pull-down assays. His-T7tag-PpNSE5(aa1-230), His-T7tag-PpNSE5(aa72-230) constructs and pET-28c(+) empty vector were expressed in bacteria, prebound on anti-T7tag agarose beads, and then *in vitro* radiolabelled PpSMC5(aa201-890) or PpSMC6(aa226-510+654-955) protein was applied. While PpNSE5(aa1-230) bound both PpSMC subunits (lane 3), PpNSE5(aa72-230) bound only PpSMC5 but not PpSMC6 (lane 6), suggesting the essential role of the first 71 amino acids for PpNSE5-PpSMC6 interaction. The bacterially expressed His-T7tag-PpNSE5 bait proteins were analysed using the anti-T7 antibody. The other details of pull-down and Y2H assays are as in Fig. 1.

To see whether the PpNSE5 alternative Met72 start could affect this interaction, we also prepared His-T7tag-PpNSE5(aa72-230) construct and tested its binding to PpSMC5 and PpSMC6 in the pull-down assay (Fig. 2B). Interestingly, this PpNSE5(aa72-230) fragment was able to interact with PpSMC5, but not with the PpSMC6 subunit (lane 6), suggesting that N-terminal 71 amino acids are essential for interaction with PpSMC6 but not PpSMC5.

### PpSMC6 joint region is required for PpNSE5 and PpNSE6 binding

To analyse PpNSE5-PpSMC6 interaction further, we prepared different arm fragments of PpSMC6 truncated from the head-or hinge-proximal end (Fig. 3A; (Lelkes *et al.*, 2023)). The long headless fragments Gal4BD-PpSMC6(aa226-955) and (aa255-923) bound both Gal4AD-PpNSE5(aa1-230) and Gal4AD-PpNSE6(aa1-370) (Fig. 3B, lanes 1 and 2). In contrast, the shorter headless fragment Gal4BD-PpSMC6(aa290-870) failed to interact with PpNSE5 and PpNSE6 subunits (Fig. 3B, lane 3). Notably, these fragments showed the functional ability to bind the PpSMC5 (aa360-710) fragment containing the hinge region (Fig. S2A; (Sergeant *et al.*, 2005, Alt *et al.*, 2017)). These data suggest the essential role of the PpSMC6 joint region (aa255-290 and aa870-923) for PpNSE5 and PpNSE6 binding.

**Figure 3.**
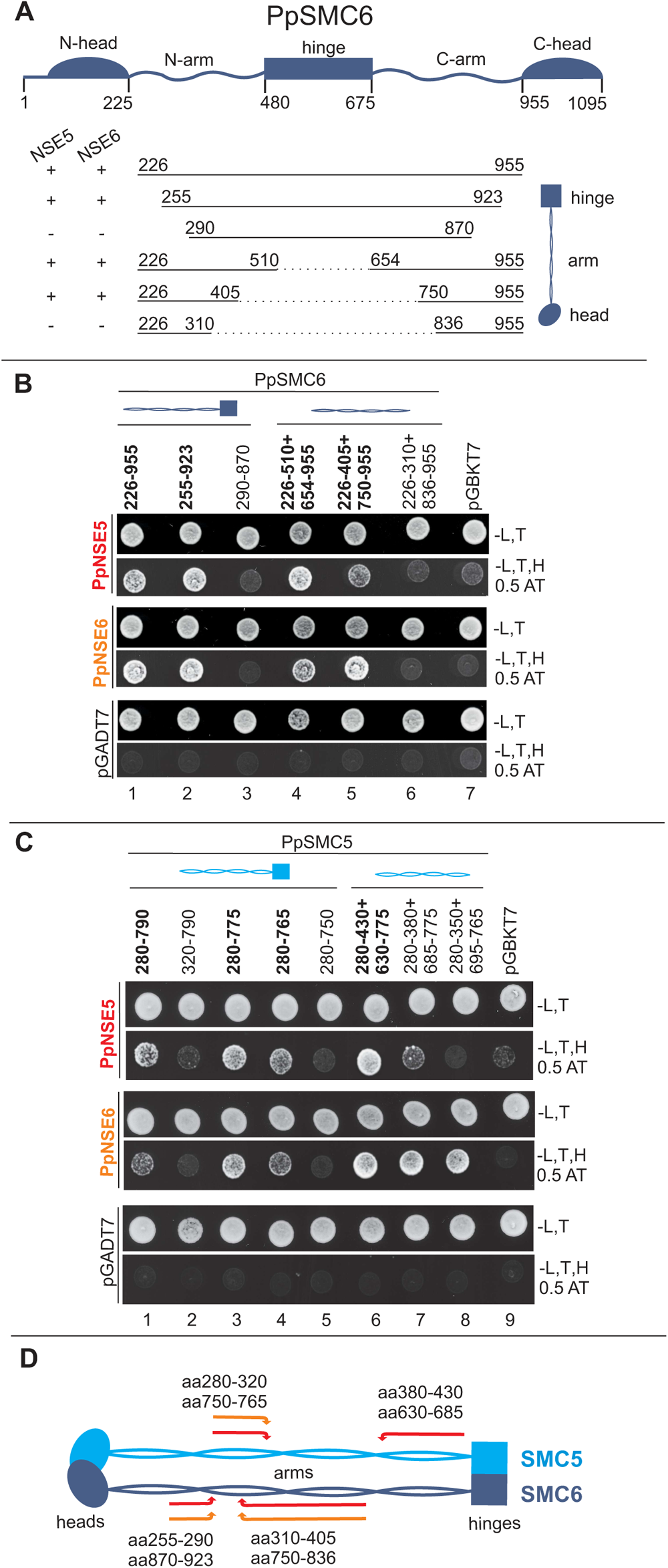
Detailed analysis of PpSMC5 and PpSMC6 interactions with PpNSE5. **(A)** Schematic representation of the PpSMC6 polypeptide chain, containing N- and C-terminal parts of the head domain, N- and C-terminal helical segments of the arm, and the hinge domain. The polypeptide folds back at the hinge domain, and its N-/C-terminal parts assemble to the coiled-coil arm and ATPase head (vertical scheme on the right). Y2H results from PpSMC6 binding analysis to PpNSE5 and PpNSE6 (panel B) are summarized on the left: +, binding; –, not binding. **(B)** Gal4BD-PpSMC6 constructs (depicted in panel A) were analysed for their binding to Gal4AD-PpNSE5(aa1-230) (top panels) and Gal4AD-PpNSE6(aa1-370) (middle panels). The Gal4BD-PpSMC6(aa226-955) and (aa255-923) fragments were able to bind PpNSE5 and PpNSE6 constructs, while the Gal4BD-PpSMC6(aa290-870) fragment was not (lanes 1-3), suggesting an essential role of the joint region for these interactions. Similarly, Gal4BD-PpSMC6(aa226-510+654-955) and Gal4BD-PpSMC6(aa226-405+750-955) constructs bound both PpNSE5 and PpNSE6 fragments, while the Gal4BD-PpSMC6(aa226-310+836-955) construct did not. **(C)** Detailed mapping of PpSMC5 binding to PpNSE5. The Gal4AD-PpNSE5(aa1-230) and Gal4AD-PpNSE6(aa1-370) constructs bound Gal4BD-PpSMC5(aa280-790) headless construct but not its N- and C-terminally truncated versions, suggesting PpNSE5 binding similar to PpNSE6 (lanes 1-5). However, the PpNSE5 binding pattern with PpSMC5 hingeless constructs was different (lanes 6-8), suggesting that PpNSE5 occupies the hinge-proximal half of the PpSMC5 arm (aa280-430 and aa630-775), while PpNSE6 binds only its middle part (aa280-350 and 695-765). All protein-protein interactions were scored in the same way as in Fig. 1. **(D)** Summary of protein-protein interaction mapping. Arrows delineate the areas essential for PpSMC5 (light blue) and PpSMC6 (dark blue) interactions with PpNSE5 (red) and PpNSE6 (orange).

Next, we prepared PpSMC6 arm-only constructs, missing both hinge and head domains, and tested them for binding to the PpNSE5 and PpNSE6 subunits. The long constructs Gal4BD-PpSMC6(aa226-510+654-955) and (aa226-405+750-955) bound both PpNSE subunits (Fig. 3B, lanes 4 and 5), suggesting no role of head and hinge domains in PpSMC6 binding to PpNSE partners. In contrast, the short arm construct (aa226-310+836-955) was not able to bind PpNSE5 or PpNSE6 (Fig. 3B, lane 6). These data suggest that PpNSE5 and PpNSE6 require PpSMC6 sequence within aa310-405 and aa750-836 and joint region (summarized in Fig. 3D).

### The hinge-proximal part of the PpSMC5 arm is essential for PpNSE5 binding

Like with PpSMC6, we wanted to delineate the PpNSE5-binding region of PpSMC5 and used a similarly designed panel of constructs (Fig. S3A; (Lelkes *et al.*, 2023)). We found that the Gal4BD-PpSMC5(aa280-790) headless fragment interacts with both Gal4AD-PpNSE5 and Gal4AD-PpNSE6 (Fig. 3C, lane 1). These interactions were abrogated by 40 amino acid truncations either from N- or C-terminus (fragments aa320-790 and aa280-750; Fig. 3C, lanes 2 and 5), suggesting PpNSE5 binding similar to PpNSE6 (as well as to PpNSE2; Fig. S3B; (Lelkes *et al.*, 2023)). However, when we employed PpSMC5 hingeless constructs, we found different binding patterns for PpNSE5. While the longest arm construct (aa280-430+630-775) interacted with all three partners (PpNSE5, PpNSE6, and PpNSE2; Figs. 3C and S3B, lane 6), shorter hingeless constructs (aa280-380+685-775 and aa280-350+695-765) bound only PpNSE6 and PpNSE2 but not PpNSE5 (Figs. 3C and S3B, lanes 7 and 8). These results suggest that PpNSE5 requires a hinge-proximal half of the PpSMC5 arm (Fig. 3D), while PpNSE2 and PpNSE6 occupy only its middle part (aa280-350 and aa695-765).

### Generation and analysis of the moss *Ppnse5* mutants

To characterize the PpNSE5 function, we created two distinct *Ppnse5* moss mutant lines. These mutants involved the insertion of a STOP codon via CRISPR/Cas9 homology-directed repair at either amino acid position 14 (*Ppnse5KO1*) or 168 (*Ppnse5KO2*) within the *PpNSE5* open-reading frame (Fig. 4A). We successfully obtained viable mutant lines named *Ppnse5KO1-16* and *Ppnse5KO2-14*, suggesting a non-essential role of the *PpNSE5* gene similar to *PpNSE6* (Lelkes *et al.*, 2023)*. Ppnse5KO1-16* exhibited a growth rate similar to WT, while the *Ppnse5KO2-14* mutant displayed a growth rate reduction of 37% (Fig. S4A). Both mutant lines showed developmental abnormalities (Fig. 4B and C) with inhibited gametophore formation. Notably, the gametophores of *Ppnse5KO2-14* were more defective, with fewer phyllids than *Ppnse5KO1-16*.

**Figure 4.**
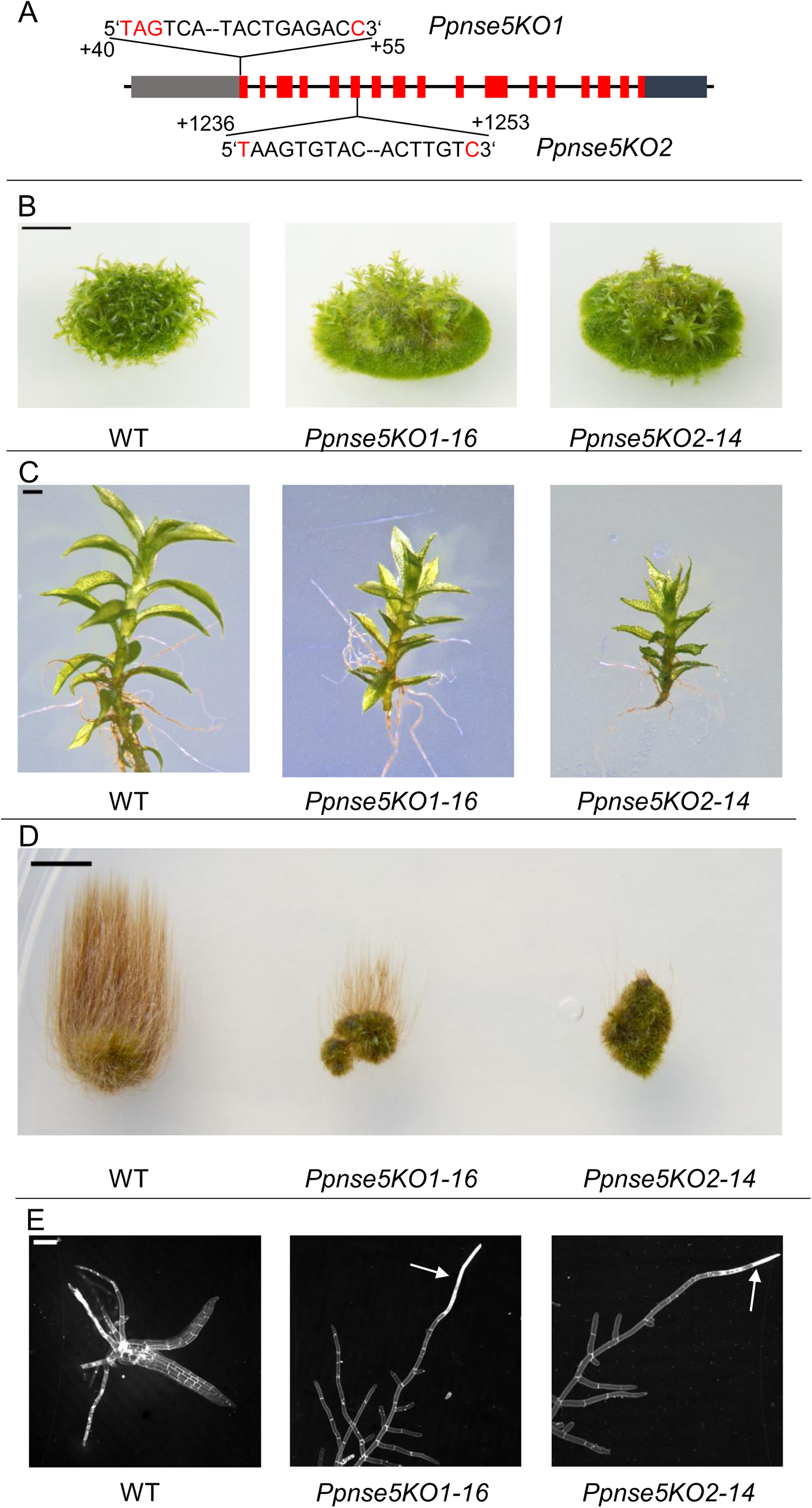
Characterization of the moss *Ppnse5* mutant lines. **(A)** Schematic representation of exon/intron distribution of *PpNSE5* gene. Detailed positions and sequences of the knock-in mutations introduced by the CRISPR/Cas9 system to generate *Ppnse5KO1* and *Ppnse5KO2* STOP-codons at aa14 and aa168, respectively. **(B-D)** Representative morphologies of *Ppnse5KO1-16* and *Ppnse5KO2-14* mutant lines suggest developmental defects in *Ppnse5* mutants. **(B)** 1-month-old colonies of WT and *Ppnse5* mutant lines grown on BCDAT medium. Scale bar = 5 mm. **(C)** Detail photos of individual gametophores of WT and *Ppnse5* mutant lines. Scale bar = 1 mm. **(D)** Comparison of caulonema development in WT and mutant lines after 3 weeks under stimulating conditions. Scale bar = 5 mm. **(E)** 10-day-old protonema stained with propidium iodide. In the WT sample, juvenile gametophores are evident, whereas the mutant lines continue to develop apical caulonemal cells, which frequently exhibit damage. Scale bar = 100 µm.

We took advantage of the development of gametophores and caulonemata (but not chloronemata) without light (Cove *et al.*, 1978) and investigated the transition from chloronema to caulonema in mutant lines. After three weeks under dark conditions, WT plants produced long, negatively gravitropic caulonemata (Fig. 4D). In contrast, both *Ppnse5KO* mutant lines exhibited a severe inhibition in the production of caulonemal filaments. Furthermore, propidium iodide (PI) staining of 10-day-old protonema revealed that the rarely occurring apical caulonemal cells were often defective, displaying premature senescence and leading to cell death in mutant lines (Fig. 4E). Microscopic analysis also showed a marked delay in bud development and, consequently, gametophore formation in both mutant lines. By day 10 after planting, WT plants had already initiated the development of juvenile gametophores. In contrast, the mutant lines had only produced apical caulonemal cells, suggesting the role of PpNSE5 in moss development. Interestingly, the *Ppnse5KO2-14* phenotypes were pronounced more than in *Ppnse5KO1-16* (Fig. 4B-D).

### Role of *PpNSE5* in the maintenance of genome stability

Based on the best-known role of SMC5/6 subunits in DNA repair and replication (Aragón, 2018, Palecek, 2019), we determined the growth response of the *Ppnse5* mutant lines to exogenous DNA damage (Holá *et al.*, 2021). As expected, both lines were sensitive to the radiomimetic drug bleomycin, although to a different extent (Fig. 5A). While growth activity in WT decreased to 77% after treatment with 30 µg BLM, growth in *Ppnse5KO1-16* and *Ppnse5KO2-14* was reduced to 63% and 35%, respectively.

**Figure 5.**
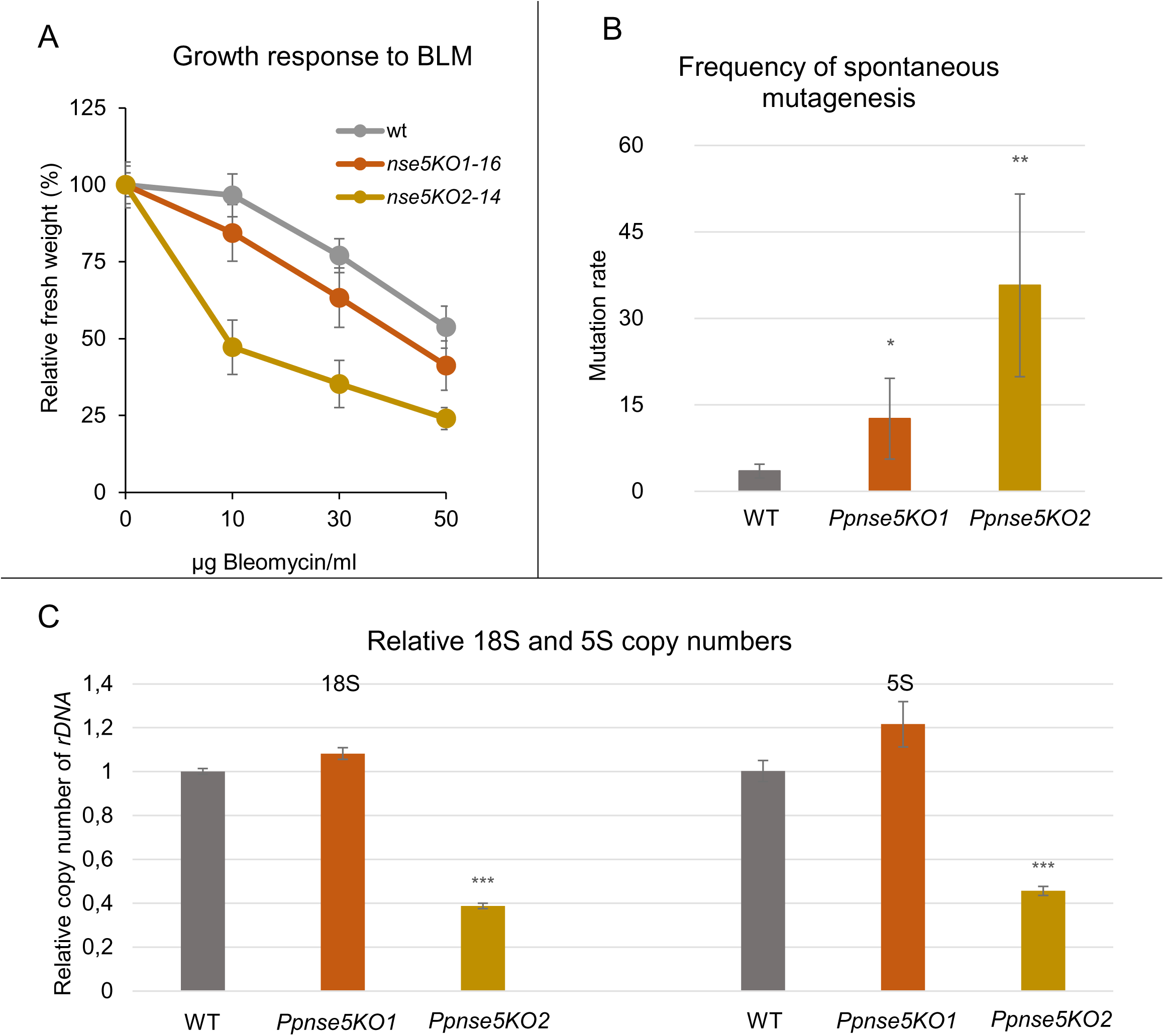
Analysis of the genome stability maintenance in *Ppnse5* mutant lines. **(A)** The growth response of WT and *Ppnse5* plants treated for 1 hr with 10, 30, and 50 µg/ml bleomycin (BLM). The explants were inoculated on a BCDAT medium and grown under standard conditions for three weeks. The mean weight of treated explants was normalized to the weight of untreated explants (set as a default 100%). *Ppnse5* mutants exhibited reduced DNA repair efficiency. **(B)** The spontaneous mutations in the *APT* gene lead to resistance to 2-FA. The 2-FA surviving colonies were counted and expressed as the number of 2-FA resistant colonies per mg of dry tissue. Spontaneous mutagenesis is increased to 12.6 in *Ppnse5KO1-16* and 35.7 in *Ppnse5KO2-14* vs. only 3.5 colonies per mg of the dry tissue in the WT. **(C)** The relative number of *18S* and *5S rDNA* copies was measured by qPCR in WT (default set to 1) *Ppnse5* mutant lines. Surprisingly, the *Ppnse5KO1-16* mutant exhibited WT-like levels of rDNA copies. Student’s t-test: \**P* < 0.05; \*\**P* < 0.01; \*\*\**P*<0.001, and error bars represent SD between the means of biological replicates.

In addition, we analysed spontaneous mutagenesis using the reporter system that depends on inactivation of the adenine phosphoribosyl transferase (*APT*) gene and leads to resistance to halogenated 2-fluoro adenine (2-FA) base (Trouiller *et al.*, 2006). Spontaneous *APT* mutations manifest as increased numbers of 2-FA resistant plants regenerated from *Ppnse5* mutant lines. While there were 3.5 resistant colonies per mg of dry tissue in the WT, it increased to 12.6 in *Ppnse5KO1-16* and 35.7 in *Ppnse5KO2-14* (Fig. 5B), suggesting a reduced capacity of *Ppnse5* mutant lines to process endogenously induced DNA damage of whatever origin. As for exogenous damage (Fig. 5A), the DNA repair efficiency of endogenous damage was reduced more in *Ppnse5KO2-14* than in *Ppnse5KO1-16* (Fig. 5B).

Given the role of SMC5/6 in *rDNA* stability maintenance (Torres-Rosell *et al.*, 2007, Peng *et al.*, 2018, Lelkes *et al.*, 2023), we measured changes in the number of *rDNA* copies in genomic DNA of 7-day-old protonemata by qPCR. Surprisingly, despite the developmental and DNA-repair problems of the *Ppnse5KO1-16* mutant, it exhibited WT-like levels of rDNA copies (Fig. 5C). In contrast, we found *rDNA* copy levels of *Ppnse5KO2-14* significantly reduced (copy numbers of *18S rDNA* were reduced to 38% and *5S rDNA* to 45%; Fig. 5C), suggesting different effects of the STOP codon insertions on different SMC5/6 functions.

The most likely explanation for the milder phenotypes of *Ppnse5KO1-16* compared to *Ppnse5KO2-14* is that the STOP codon at amino acid position 14 does not abolish PpNSE5 translation completely. We assumed that PpNSE5 translation might start at the alternative ATG codon (Met72), producing the N-terminally truncated PpNSE5(aa72–526) protein. We introduced the 3xFLAG-tag at the 3’-end of PpNSE5 ORF to test this possibility and analysed its expression in WT and KO1 lines using the anti-FLAG antibody. We observed a clear band in the WT-FLAG (FL PpNSE5) extract corresponding to its theoretical size of 62 kDa and a lower band of roughly 55 kDa in the extract from *Ppnse5KO1*-FLAG line (Fig. S4C, lanes 2 and 3), which corresponds to the PpNSE5(aa72–526)-3xFLAG translation product (theoretical size of 54 kDa). Such PpNSE5(aa72–526) protein should be able to interact with PpNSE6 and PpSMC5 partners (Figs. 1C and 2B) and therefore assemble the SMC5/6 complex. On the other hand, the inability of the PpNSE5(aa72–526) protein to bind PpSMC6 (Fig. 2B) could partially compromise SMC5/6 function and result in relatively mild phenotypes (see discussion).

## DISCUSSION

Here, we characterized PpNSE5 binding to PpNSE6 and PpSMC subunits of *P. patens* PpSMC5/6 complex. Our analyses showed the binding of PpNSE5 to the head-proximal region of PpSMC6 and the hinge-proximal arm of PpSMC5. Consistent with our PpNSE5-PpSMC6 interaction data, the crosslinking studies of budding yeast scSMC5/6 suggested the proximity of scNSE5 to scSMC5-scSMC6 heads (Gutierrez-Escribano *et al.*, 2020, Taschner *et al.*, 2021, Yu *et al.*, 2021). However, recent cryoEM structure studies narrowed down the binding site of the yeast scNSE5-scNSE6 dimer to scSMC6 neck (Li *et al.*, 2023), while our analyses localized PpNSE5 (and PpNSE6) to PpSMC6 join region (Fig. 3). As the yeast and plant NSE5 (as well as NSE6) sequences exhibit very low homologies (Oravcova *et al.*, 2022, Lelkes *et al.*, 2023), the (slightly) different NSE5-SMC6 binding modes most likely reflect evolutionary distance between yeast and plants.

Nevertheless, these different binding modes may still serve similar functions, given another interesting comparison between binding sites of NSE5-SMC5 in yeasts and plants. It was previously reported that yeast scSMC5-scSMC6 hinge fragments co-purify with scNSE5-scNSE6 dimers (Duan *et al.*, 2009). In comparison, our results show the binding of PpNSE5 to the hinge-proximal arm of PpSMC5 (Fig. 3). Although the binding modes are again (slightly) different between yeasts and plants, the mapping of SMC5 (hinge/hinge-proximal) and SMC6 (neck/joint) sites suggests binding of NSE5 to very distant regions within the SMC5/6 complexes (Fig. 6A). The binding sites at hinge/hinge-proximal and neck/joint regions are tens of nanometres apart in rod-like structures, making it unlikely that the NSE5 molecule could bind both areas within such conformation of the SMC5/6 complex. The data rather suggest that NSE5(-NSE6 dimers) either (1) link antiparallel SMC5/6 dimers or (2) bind SMC5/6 monomers in elbow-bent conformation (Fig. 6A).

**Figure 6.**
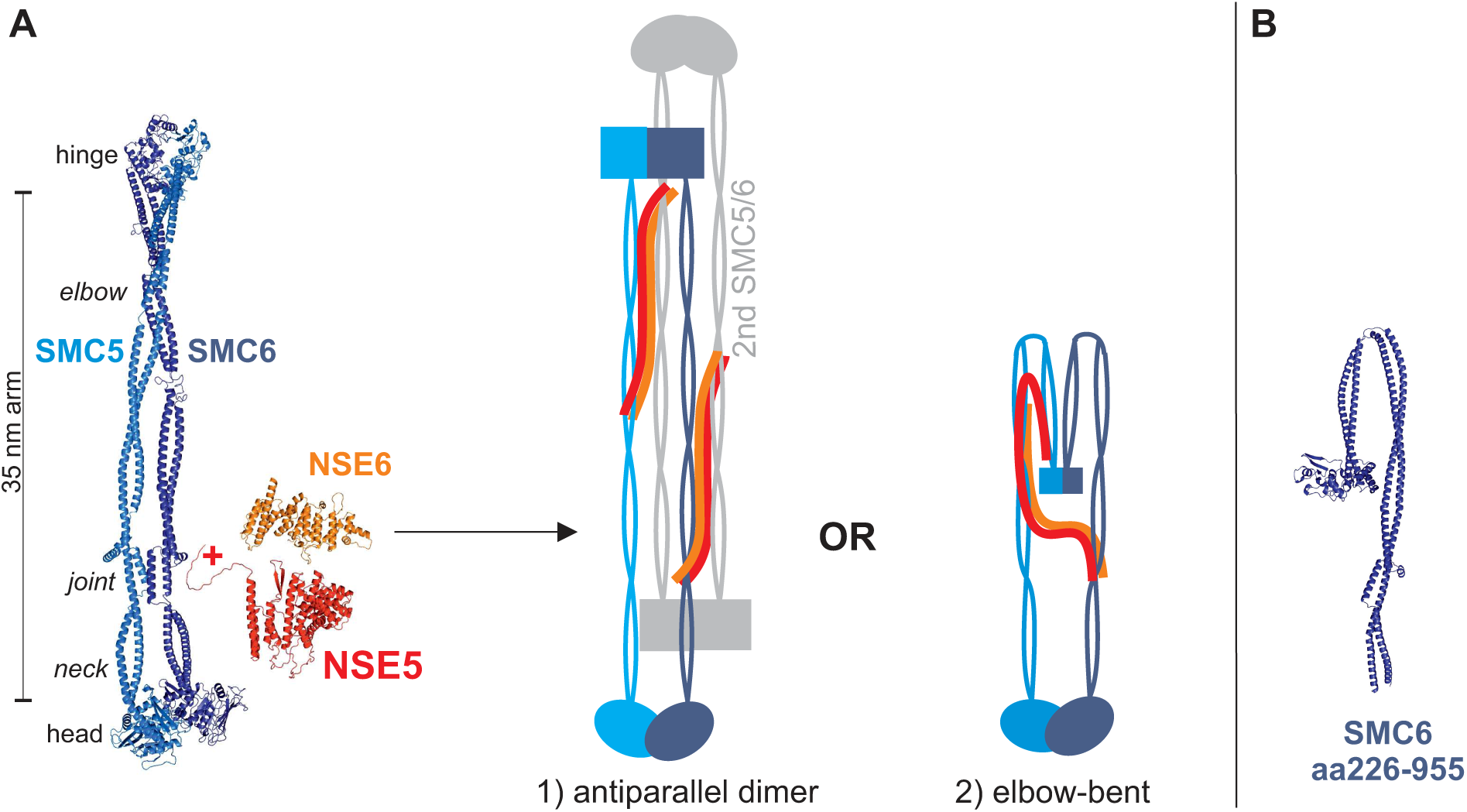
Hypothetical models for NSE5/NSE6 binding to distant regions of the SMC arms. **(A)** Structural rod-like model of PpSMC5-PpSMC6 dimer (based on the 7QCD cryoEM structure; (Hallett *et al.*, 2022)) compared to the PpNSE5 (red) and PpNSE6 (orange) AlphaFold models. The distance between the head and hinge domains set by the long coiled-coil arm is approximately 35 nm, making it unlikely that mostly globular small NSE5 molecule could bind both the SMC6 joint region and SMC5 hinge-proximal end at the same time. We hypothesize that either 1) the NSE5 molecule binds the SMC6 joint region of the first SMC5-SMC6 dimer (light and dark blue colours, respectively) and SMC5 hinge-proximal end of the second antiparallel SMC5-SMC6 dimer (grey colours) in rod-like conformation or 2) SMC5-SMC6 dimer bends at the elbow and the hinges come closer to the heads, allowing SMC6-joint-bound NSE5 to reach SMC5 hinge-proximal end. The other NSE subunits were omitted for simplicity. **(B)** The AlphaFold model of the PpSMC6(aa226-955) headless fragment has a bent arm at the elbow region.

Interestingly, dimers of SMC5/6 complexes (model 1; Fig. 6A) were recently implicated in loop extrusion (Pradhan *et al.*, 2023). However, NSE5-NSE6 subunits rather blocked the formation of productive dimers and inhibited loop extrusion. It is possible that loop extrusion requires parallel SMC5/6 dimers while NSE5-NSE6 drives the formation of antiparallel dimers, which are probably incompatible with the loop extrusion. However, given no experimental evidence of NSE5/NSE6-containing SMC5/6 dimers so far, we favour the elbow-bent model (model 2; Fig. 6A). In fact, this model is supported by low-resolution cryoEM structures of scSMC5/6 (co-purified with scNSE5-scNSE6 subunits), showing elbow-bent conformations of the complexes (Gutierrez-Escribano *et al.*, 2020). In addition, our AlphaFold modelling of PpSMC5 and PpSMC6 headless fragments provided the elbow-bent structures (Fig. 6B and not shown) reminiscent of cohesin and condensin (Lee *et al.*, 2020, Petela *et al.*, 2021). As the bending of cohesin and condensin complexes plays a key role in their loading to DNA (Bürmann and Löwe, 2023), it is reasonable to assume that the binding of the NSE5-NSE6 dimer to bent SMC5/6 assists with its loading. Indeed, the NSE5-NSE6 dimer is considered the major loading factor of SMC5/6 to DNA lesions (Oravcová and Boddy, 2019).

Our *in vivo* results are consistent with the above NSE5-NSE6 loading function and *in vitro* data. We showed that the first 71 amino acids of PpNSE5 are specifically required for binding to PpSMC6 while dispensable for interactions with PpNSE6 and PpSMC5 (Figs. 1 and 2). Interestingly, the deletion of the first 71 amino acids of PpNSE5 in *P. patens* plants (*Ppnse5KO1-16* mutant line) resulted in compromised DNA repair functions (and milder developmental defects; Fig. 4) while not affecting rDNA copy numbers (Fig. 5). These results suggest that PpNSE5-PpSMC6 interaction is specifically involved in DNA repair-related functions of PpSMC5/6 but not in its replication-coupled activities. This is consistent with NSE5-NSE6 function as the SMC5/6 loader to DNA lessions found in yeasts and humans (Räschle *et al.*, 2015, Oravcová and Boddy, 2019). Strikingly, a recent study showed that yeast mutant cells defective in scNSE6-scSMC6 interaction are sensitive to DNA damage but do not exhibit replication-coupled problems (Li *et al.*, 2023). In conclusion, the binding of NSE5-NSE6 dimer to SMC6 plays a specific role in DNA repair, and this function seems to be conserved (at least from yeast to plants). It will be interesting to reexamine human hsNSE5 binding to hsSMC5 and hsSMC6 subunits (Adamus *et al.*, 2020), and compare their binding sites to yeasts and plants. Altogether, our detailed analysis has already provided new insights into NSE5 features conserved across species that were previously assumed not to be conserved.

## METHODS

### Cloning of constructs

For the Y2H constructs, the full-length PpNSE5(aa1-526) construct created in (Lelkes *et al.*, 2023) was used as a template to create shorter PpNSE5 fragments (Figs. 1 and 2). Individual fragments were amplified using specific primers listed in Supp. Table ST1. PCR products were cloned into the *Nde*I-*Bam*HI sites of pGBKT7 or pGADT7 vectors using NEBuilder (New England BioLabs, USA).

The cloning of pGADT7-PpNSE6 (full length and aa1-370), pGADT7-PpNSE2(aa1-304), pGBKT7-PpSMC5 (containing aa280-790, aa320-790, aa280-775, aa280-765, aa280-750, aa360-710, and aa280-350+695-765), and pGBKT7-PpSMC6 (containing aa226-955, aa290-870, and aa226-510+654-955) plasmids have been described in (Lelkes *et al.*, 2023). The PpSMC6(aa255-923) fragment was amplified using BK005 and BK006 primers (Supp. Table ST1), and cloned into the *Nde*I-*Bam*HI site of pGBKT7 vector using NEBuilder to obtain Gal4BD-PpSMC6(aa255-923) construct. The Gal4BD-PpSMC5(aa280-430+630-775), Gal4BD-PpSMC5(aa280-380+685-775), Gal4BD-PpSMC6(aa226-405+750-955), and Gal4BD-PpSMC6(aa226-310+836-955) hingeless constructs (Fig. 3) were cloned in two steps. First, the N-terminal part was amplified by PCR and cloned into the pGBKT7 *Nde*I site. Second, the C-terminal part was PCR amplified and inserted into the *Bam*HI site of the N-terminal construct created in the first step (Supp. Table ST1).

For the pull-down assays, pET-Duet-PpNSE6(aa1-370)-StrepTwin-Stag and pET-28a-PpNSE5(aa1-526) plasmids with codon optimized for *E. coli* expression were described in (Lelkes *et al.*, 2023). The PpNSE5 fragments aa1-230 and aa72-230 (Fig. 2B) were amplified using the primers listed in the Supp. Table ST1. Purified inserts were cloned into the pET-28c(+) vector containing His-T7tag (*Bam*HI-*Xho*I) using NEBuilder.

### Yeast two-hybrid assay

The classical Gal4-based Y2H system was used to analyse protein-protein interactions as described previously (Hudson *et al.*, 2011). Briefly, pGBKT7- and pGADT7-based plasmids were co-transformed into the *S. cerevisiae* PJ69–4a strain and selected on SD -Leu, -Trp plates. Drop tests were carried out on SD -Leu, -Trp, -His (with 0.1; 0.2; 0.3; 0.5; 1; 2; 3; 4; 5; 10; 15; 20; 30 mM 3-aminotriazole) plates at 28°C. Each combination was co-transformed at least three times, and three independent drop tests were carried out.

### Pull-down assays

In a pull-down analysis examining the interaction of the PpNSE6 and PpNSE5 subunits (Fig. 1D), T7-driven *in vitro* expression from plasmids pGBKT7-PpNSE5(aa190-526), pGBKT7-PpNSE5(aa245-526), and pET-Duet-PpNSE6(aa1-370)-StrepTwin-Stag was performed using the TNT kit (Promega, USA). Radiolabeled methionines (Met S^35^, Hartmann Analytic, Germany) were incorporated into the sequences of the synthesized proteins. Then, PpNSE5 proteins were mixed with PpNSE6 and preincubated at 4°C for 0.5 h in a total volume of 300 ul 1× phosphate buffer (10× phosphate buffer consisting of 42.9 mM Na_2_HPO_4_, 14.7 mM KH_2_PO_4_, 27 mM KCl, 1.37 M NaCl, 1% Tween-20, 0.02% sodium azide, pH 7.3). Mix was added to the protein-S beads and incubated for 1.5 hours at 4°C. Input, unbound, and bound fractions were separated by 12% SDS-PAGE, transferred to nitrocellulose membranes, and analysed by phosphorimaging.

For the His-T7tag-PpNSE5 pull downs (Fig. 2B), constructs were transformed to BL21(DE3)RIL strain. Transformed bacteria were further inoculated and grown in liquid LB medium at 37 °C to early exponential phase (OD_600_ ∼ 0.5). Expression was induced with 0.5 mM IPTG, and cultures were incubated at 37°C for 5 hours. The harvested cells were resuspended in 1× phosphate buffer and sonicated with constant cooling on ice. Cleared lysates were incubated for 2 hours on anti-T7-tag agarose beads and washed three times with 1× phosphate buffer. Then, Met-S^35^ radiolabeled proteins [PpSMC5(aa201-890) or PpSMC6(aa226-510+654-955)] expressed *in vitro* were added and incubated overnight at 4°C in a total volume of 200 μl of 1x phosphate buffer. Input, unbound, and bound fractions were separated by 12% SDS-PAGE, transferred to nitrocellulose membranes, and analysed by phosphorimaging and immunoblotting with the anti-T7tag HRP antibody (Abcam - ab19291, USA).

### *In silico* protein analysis

The PpNSE5 and PpNSE6 AlphaFold models from the EMBL-EBI database were used (Varadi *et al.*, 2022). The rod-like PpSMC5 and PpSMC6 structural models were generated with the Swiss-Model tool using 7QCD cryoEM structures as templates (Schwede *et al.*, 2003, Hallett *et al.*, 2022). The headless PpSMC5 and PpSMC6 fragments were modelled using the AlphaFold/ColabFold tool (Mirdita *et al.*, 2022). Secondary structure and fragment borders were determined based on the 3D models. The structures were aligned and visualized using the PyMOL software (Schrodinger Inc., USA).

### Plant material cultivation and analysis

The wild-type (WT) *P. patens*, accession (Hedw.) B.S.G. (Rensing *et al.*, 2008), was utilized to create the *Ppnse5* mutants. The moss lines were cultivated either as ’spot inocula’ on BCD agar medium enriched with 1 mM CaCl_2_ and 5 mM ammonium tartrate (BCDAT medium) or as lawns of protonema filaments by subculturing homogenized tissue on BCDAT agar, overlaid with cellophane, within growth chambers maintained at an 18/6-hour day/night cycle and a temperature of 22/18°C (Cove *et al.*, 2009). In dark conditions, 0.5% sucrose was included in the medium, and 1.5% agar was used. Petri dishes were positioned vertically to enhance the observation of caulonema growth.

The growth rates were determined by weighing untreated explants of mutant lines and comparing them with untreated explants of WT. Sensitivity to DNA damage was measured as described previously (Lelkes *et al.*, 2023). Protocols for DNA isolation and rDNA copy numbers analysis were also detailed in (Lelkes *et al.*, 2023).

The pictures of whole colonies were taken by Canon EOS77D, objective Canon EF 28-135mm f/3.5-5.6. The details of gametophores were photographed by Stereomicroscope Leica M205FA, objective Plan-Apochromat 2.0x. Staining with 10 µg/ml propidium iodide (PI, Sigma-Aldrich) was performed as described previously (Lelkes *et al.*, 2023).

### Construction of *Ppnse5* mutant lines

The STOP codon knock-in was achieved through homology-directed repair following the induction of double-strand breaks (DSB) within the *PpNSE5* locus by Cas9. We used two double-stranded DNA donor templates, one 45 bp and the other 40 bp in length, to introduce the desired mutations for constructing *Ppnse5KO1* and *Ppnse5KO2*, respectively. These donor templates were synthesized as pairs of complementary oligonucleotides (pKA1390 and pKA1391 for *Ppnse5KO1*; pKA1394 and pKA1395 for *Ppnse5KO2*; see Supp. Table ST2) designed to be homologous to the first and fifth exons of the *PpNSE5* locus (Fig. 4A). The *Ppnse5KO1* template contained a +40CCT to TAG substitution (resulting in a STOP codon at the 14th amino acid position) and a +55A to C substitution, which generated a *Bsa*I cleavage site. The *Ppnse5KO2* template featured a +1236A to T substitution (resulting in a STOP codon at the 168th amino acid position) and a +1253A to C substitution, creating a *Sal*I cleavage site.

We utilized the Gateway destination vector pMK-Cas9 (Addgene plasmid #113743) containing the Cas9 expression cassette and kanamycin resistance marker, along with the entry vector pENTR-PpU6sgRNA-L1L2 (Addgene plasmid #113735), which harboured the PpU6 promoter and sgRNA scaffold, as previously described (Mallett *et al.*, 2019). PpNSE5-specific sgRNA spacers were synthesized as pairs of complementary oligonucleotides (pKA1392 + pKA1393 for *Ppnse5KO1* and pKA1396 + pKA1397 for *Ppnse5KO2*; see Supp. Table ST2). We added four nucleotides to the 5′-ends of these oligonucleotides to create sticky ends compatible with *Bsa*I-linearized pENTR-PpU6sgRNA-L1L2. The Cas9/sgRNA expression vectors were generated using the Gateway LR reaction, recombining the entry vector with the *PpNSE5* sgRNA spacers and the destination vector.

These DNA constructs, together with the appropriate donor templates (annealed pKA1390 + pKA1391 for *Ppnse5KO1* or pKA1394 + pKA1395 for *Ppnse5KO2*), were co-transformed into protoplasts via PEG-mediated transformation. All sgRNAs were designed in the CRISPOR online software using *P. patens* (Phytozome V11) and *S. pyogenes* (5′ NGG 3′) as the genome and PAM parameters, respectively. The protospacers with the highest specificity score were chosen.

After five days of regeneration, the transformed protoplasts were transferred to the BCDAT medium supplemented with 50 μg/l G418 for selection. Following one week of selection, the G418-resistant lines were propagated. Crude extracts from young tissues of these lines were used for PCR amplification of genomic DNA surrounding the editing sites, employing primers pKA1437 + pKA1438 for *Ppnse5KO1* and pKA1439 + pKA1440 for *Ppnse5KO2*. The resulting PCR products were cleaved using *Bsa*I or *Sal*I and subjected to DNA electrophoresis. Lines for which the PCR product was successfully cleaved underwent sequencing to confirm the accurate introduction of mutations.

We identified two lines, *Ppnse5KO1-16* and *Ppnse5KO1-23*, with correctly integrated STOP codons in the first exon and two lines, *Ppnse5KO2-14* and *Ppnse5KO2-21*, with accurately incorporated STOP codons in the fifth exon. Both mutants of the respective mutation exhibited similar morphology and growth responses to bleomycin treatment. We chose the *Ppnse5KO1-16* and *Ppnse5KO2-14* lines for further experiments with a slightly more pronounced phenotype.

### Evaluation of the frequency of spontaneous mutations

The frequency of spontaneous mutations was assessed using the adenine phosphoribosyl transferase (*APT*) gene reporter system (Trouiller *et al.*, 2006). Loss of function in the *APT* gene confers resistance to a toxic adenine analogue known as 2-fluoro adenine (2-FA). To evaluate this phenotype, tissue samples from 7-day-old wild-type (WT) and mutant lines were homogenized and then incubated in 10 ml of liquid BCDAT medium for 24 hours to facilitate recovery. Subsequently, 1 ml of the homogenized tissue was used to determine the dry tissue weight, while the remaining tissue was planted onto BCDAT medium supplemented with 5 µM 2-FA. After three weeks of incubation, the number of the 2-FA resistant colonies was counted. The frequency of spontaneous mutagenesis is given as the number of 2-FA resistant colonies per mg of dry tissue. The experiments were repeated four times.

## AUTHOR CONTRIBUTIONS

Conceptualization, Supervision, and Funding Acquisition: J.J.P., K.J.A., B.Š.; Data Curation and Formal Analysis: J.J.P., J.V., E.L., K.J.A., M.H.; Investigation: J.V., M.H., E.L.,

R.V., B.Š., B.K.; Visualization: J.J.P., J.V., K.J.A., M.H.; Writing – Original Draft Preparation: J.J.P., K.J.A., M.H., J.V.

## ACKNOWLEDGEMENTS

We acknowledge the Imaging Facility of the Institute of Experimental Botany AS CR, supported by the MEYS CR (LM2018129 Czech-BioImaging).

