## Supporting Information for "NSE5 subunit interacts with distant regions of the SMC arms in the *Physcomitrium patens* SMC5/6 complex"

#### Supporting methods

Most methods are described in the main paper.

#### Protein expression analysis

Yeast PJ69-4 cells were grown in YPD to $OD_{600}$ ~ 2.5, harvested by centrifugation, and lysed by incubation in 0.1 M NaOH for 5 min and boiling in SDS buffer (60 mM Tris–HCl, 2% SDS, 4% β-mercaptoethanol, 5% glycerol, 0.002% bromophenol blue; (Kushnirov, 2000)). Samples were separated by 12% SDS-PAGE, blotted, and stained with 0.5% Ponceau S red (Sigma-Aldrich) in 1% acetic acid. After destaining, blots were incubated with anti-Myc-HRP (Abcam - ab62928) in 1:5000 dilution for 1 h at room temperature. After washing in TBST buffer, chemiluminescence was detected using Super SignalTM West Dura Extended Duration Substrate (ThermoFisher Scientific).

The moss lines were cultivated as lawns of protonema filaments on the BCDAT agar plate. Plant material was scraped from the plate and dried on a paper towel. Protein lysates were prepared using 0.1 M NaOH, separated by 10% SDS-PAGE, and analyzed as described above (anti-FLAG-HRP, Sigma Aldrich – A8592, 1:3000 dilution).

#### RNA isolation and qRT-PCR analysis

Total RNA was isolated from 7-days-old protonemata with RNeasy Plant Mini Kit (Qiagen, USA), treated with DNaseI (DNA-free™ DNA Removal Kit, Thermo Fisher Scientific), and reverse transcribed using qPCRBIO cDNA Synthesis Kit (PCR Biosystems). Diluted cDNA reaction mixtures were used for qRT-PCR analysis using the qPCRBIO SyGreen Mix Lo-ROX (PCR Biosystems) in Stratagene-MX3005P. Analysis was performed for three biological replicas (independently cultivated tissue) and in two technical replicates with Clathrin adapter complex subunit CAP-50 (Pp3c27_2250V3.1) as a reference gene (Kamisugi *et al.*, 2016,

Lelkes *et al.*, 2023). The relative transcription of *PpNSE5* was calculated by the ΔΔCt method (Pfaffl, 2004).

**Construction of FLAG-tagged moss lines**

The FLAG-tag knock-in insertion was achieved through homology-directed repair following the induction of double-strand breaks (DSB) within the *PpNSE5* locus by Cas9. We employed the pMK-Cas9 plasmid containing PpNSE5-specific sgRNA, which was synthesized and cloned as a pair of complementary oligonucleotides (pKA1500+pKA1501). This plasmid was used to induce DSB at the 3'end of the *PpNSE5* gene, facilitating the integration of the construct with a 3xFLAG-tag linker. The 3xFLAG-tag was amplified by PCR using primers pKA1496+pKA1497, which contained sequences necessary for cloning by NEBuilder. Targeting arms flanking the 3xFLAG-tag with linker and homologous to the 3'end and 3'UTR of the *PpNSE5* gene were synthesized as pairs of complementary oligonucleotides (pKA1493 + pKA1494 and pKA1498+pKA1499), incorporating sequences for NEBuilder cloning. The resulting construct was assembled using the NEBuilder cloning system and inserted into the *EcoRI*-cleaved pBlueScript plasmid. For the transformation of protoplasts, the construct was amplified using primers pKA1502+pKA1503 and co-transformed with the Cas9/sgRNA vector. The integration of the construct was subsequently confirmed through PCR analysis using primers pKA1408+pKA1470 and verified by sequencing.

### Supporting figure legends

**Supporting Figure S1. PpNSE5 protein analysis. (A)** The unstructured N-terminal part (aa1-60) is shown as a black line; two parts of the structured region are coloured in blue (aa60-230) and red (aa230-505), respectively. PpNSE5 tertiary structure AlphaFold model (middle panel) was used to define secondary structures (bottom panel); rectangle – helix, arrow – β-sheet. **(B** and **C)** Expression of the Gal4BD-PpNSE5 constructs in yeast PJ69-4 strain (used in Figs. 1B and C) was verified using the anti-Myc-HRP antibody. The bottom panels show an equal loading of protein extracts stained with Ponceau red.

**Supporting Figure S2. PpSMC6 analysis. (A)** Indicated Gal4BD-PpSMC6 constructs were tested against Gal4AD-PpSMC5(aa360-710) fragment to verify their ability to mediate hinge-hinge interactions. The Y2H assay details as in Fig. 1. **(B)** Expression of the Gal4BD-PpSMC6 constructs in yeast PJ69-4 strain (used in Figs. 3B and S2A) was verified as in Fig. S1B.

**Supporting Figure S3. PpSMC5 analysis. (A)** Schematic representation of the different PpSMC5 constructs. Like PpSMC6 (Fig. 3A), the PpSMC5 polypeptide chain folds N- and C-terminal ends to the head domain, middle helical segments to the coiled-coil arm, and a hinge domain. Y2H results from PpNSE2, PpNSE5, and PpNSE6 binding analyses (Figs. 3C and S3B) are summarized on the right: +, binding; –, not binding. **(B)** Gal4BD-PpSMC5 constructs (panel A) were analyzed for their binding to Gal4AD-PpNSE2(aa1-304) in the same way as for PpNSE5 and PpNSE6 in Fig. 3C. PpNSE2 binds to PpSMC5 middle part (aa280-350+695-765) similar to PpNSE6 (Lelkes *et al.*, 2023). Protein-protein interactions were scored as in Fig. 1.

**(C)** Expression of the Gal4BD-PpSMC5 constructs in yeast PJ69-4 strain was verified as in Fig. S1B.

**Supporting Figure S4. Characterization of the moss *Ppnse5* mutants**. **(A)** The growth rates of WT, *Ppnse5KO1-16*, and *Ppnse5KO2-14* lines were measured as the fresh weight of 3-week-old untreated plants. **(B)** Relative levels of the *PpNSE5* transcript in the *Ppnse5KO1-16* and *Ppnse5KO2-14* mutants versus WT as measured by RT-qPCR. The quantity of cDNA was normalized against the Clathrin adapter complex subunit CAP-50. Student's t-test: ***$P$<0.001, and error bars represent SD between the means of three biological replicates. **(C)** The FLAG-tagged PpNSE5 expression. Expression from the native Met1 (FL, lane 2) produced 63.4 kDa protein, while expression from alternative Met72 resulted in 55 kDa product (KO1, lane 3, red arrow). Nonspecific bands stained by anti-FLAG antibodies are marked with an asterisk. Ponceau red staining shows equal loading of protein extracts.

**Supporting Table ST1: List of cloning primers**

**Supporting Table ST2: Primers for moss analysis and manipulation**

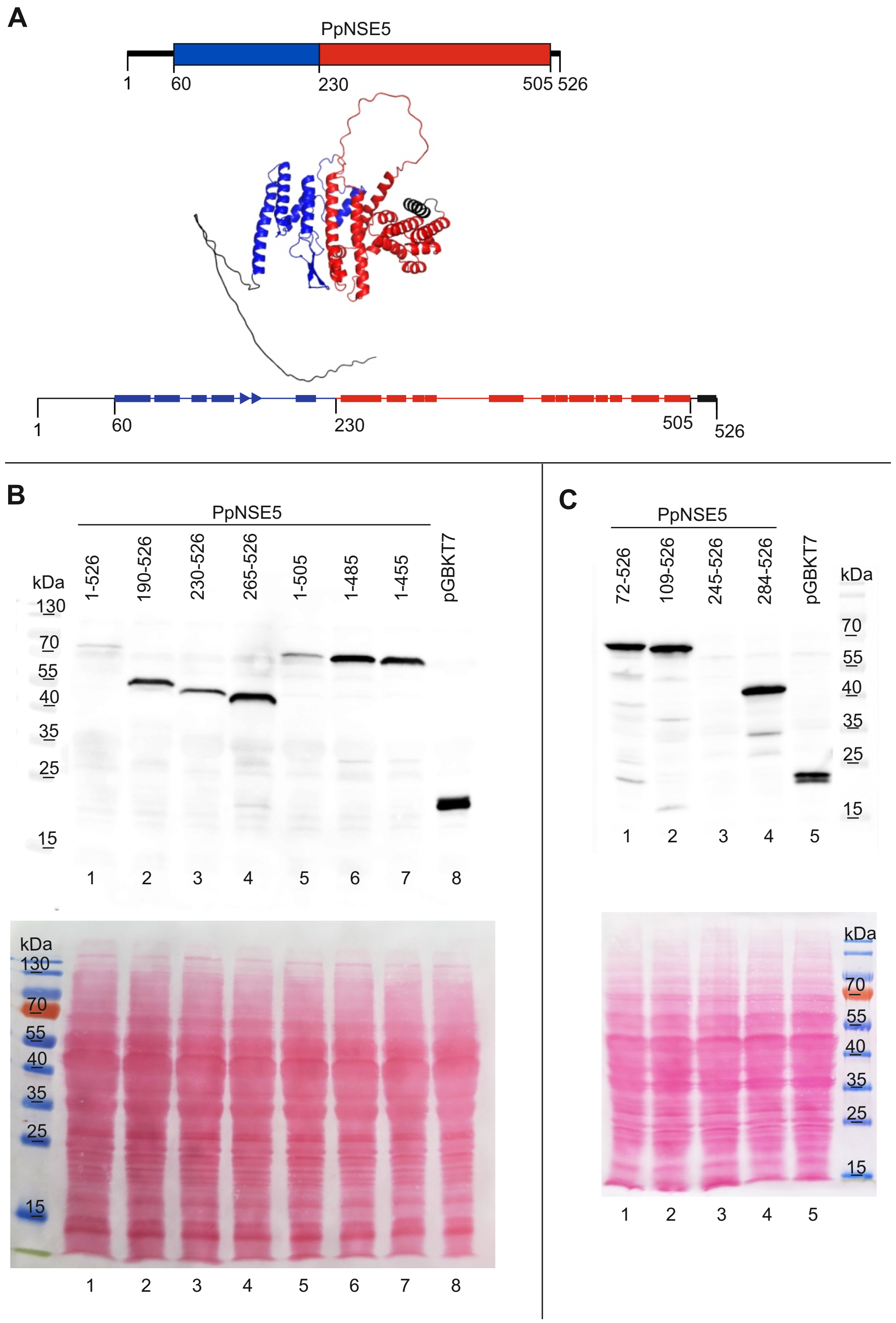
A
PpNSE5
1
60
230
505 526
1
60
230
505
526
B
PpNSE5
1-526
190-526
230-526
265-526
1-505
1-485
1-455
pGBKT7
kDa
130
70
55
40
35
25
15
1 2 3 4 5 6 7 8
kDa
130
70
55
40
35
25
15
1 2 3 4 5 6 7 8
C
PpNSE5
72-526
109-526
245-526
284-526
pGBKT7
kDa
70
55
40
35
25
15
1 2 3 4 5
kDa
70
55
40
35
25
15
1 2 3 4 5

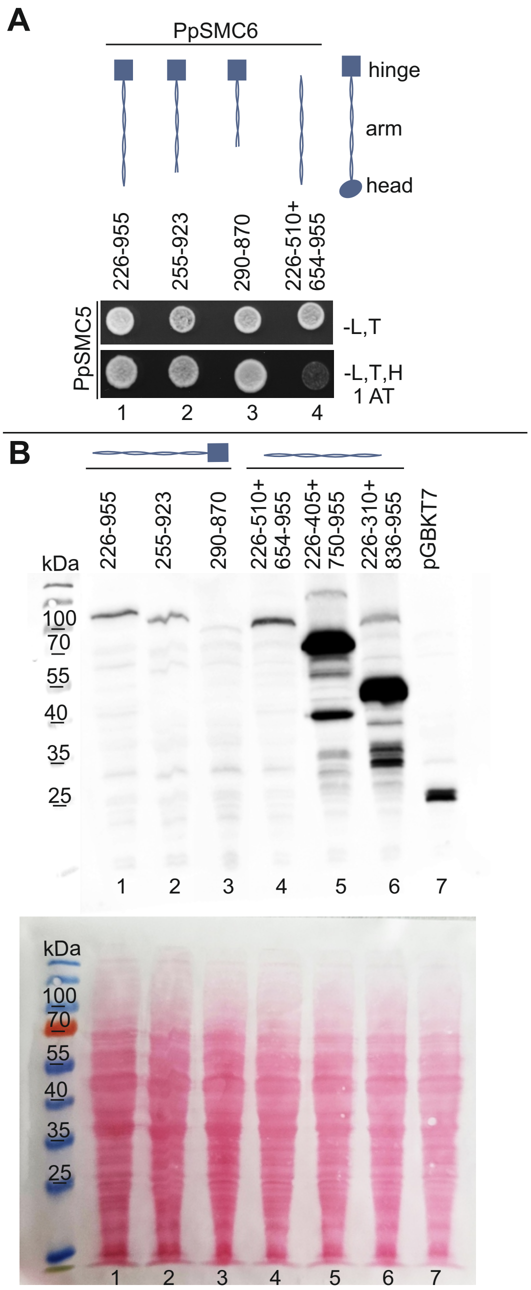
A
PpSMC6
hinge
arm
head
226-955
255-923
290-870
226-510+
654-955
PpSMC5
-L,T
-L,T,H
1 AT
1
2
3
4
B
226-955
255-923
290-870
226-510+
654-955
226-405+
750-955
226-310+
836-955
pGBKT7
kDa
100
70
55
40
35
25
1
2
3
4
5
6
7
kDa
100
70
55
40
35
25
1
2
3
4
5
6
7

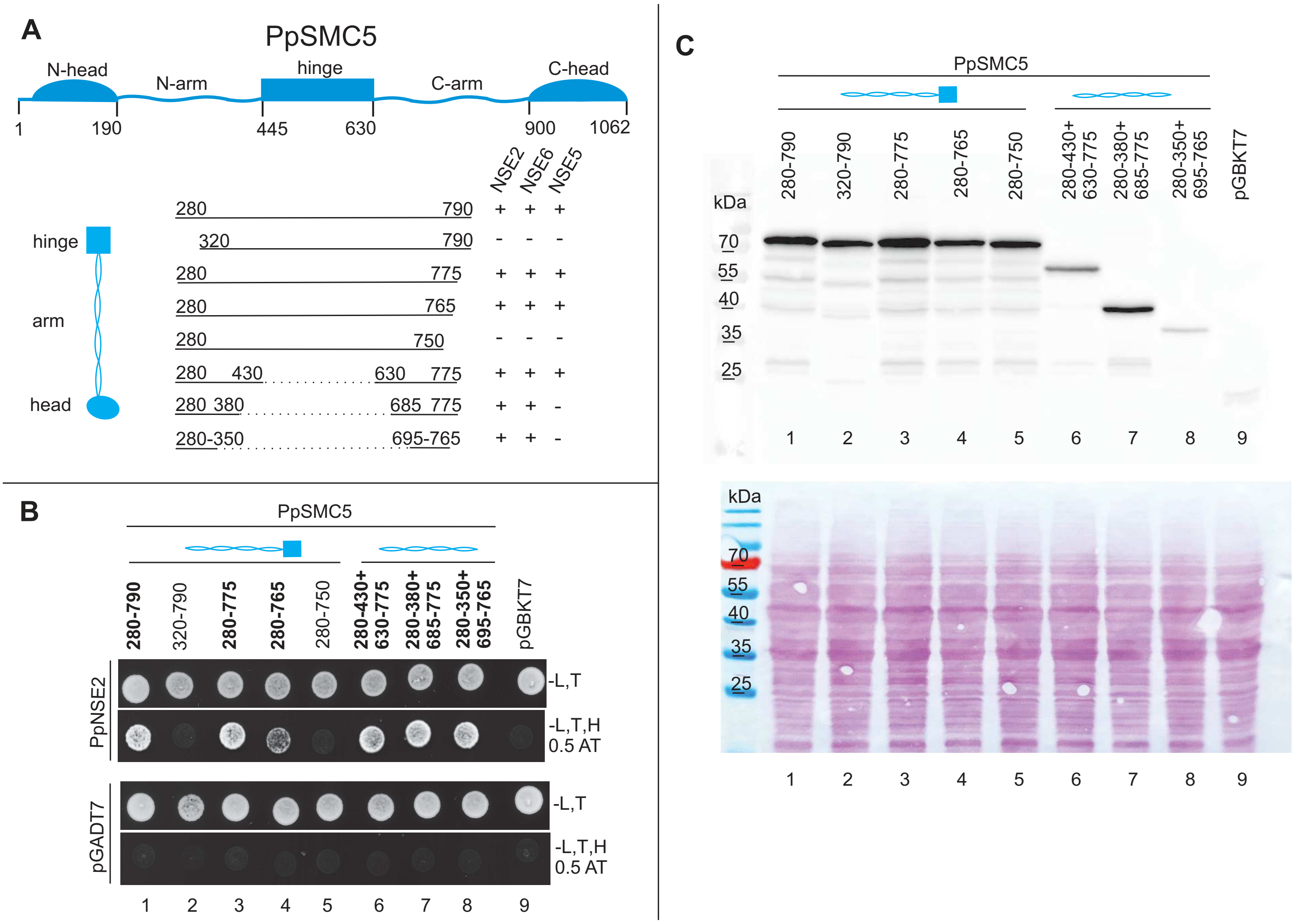

A
PpSMC5
N-head
N-arm
hinge
C-arm
C-head
1
190
445
630
900
1062
NSE2
NSE6
NSE5
hinge
arm
head
280 790 + + +
320 790 - - -
280 775 + + +
280 765 + + +
280 750 - - -
280 430 630 775 + + +
280 380 685 775 + + -
280-350 695-765 + + -
B
PpSMC5
280-790
320-790
280-775
280-765
280-750
280-430+ 630-775
280-380+ 685-775
280-350+ 695-765
pGBKT7
PpNSE2
pGADT7
-L,T
-L,T,H 0.5 AT
1 2 3 4 5 6 7 8 9
C
PpSMC5
kDa
70
55
40
35
25
1 2 3 4 5 6 7 8 9

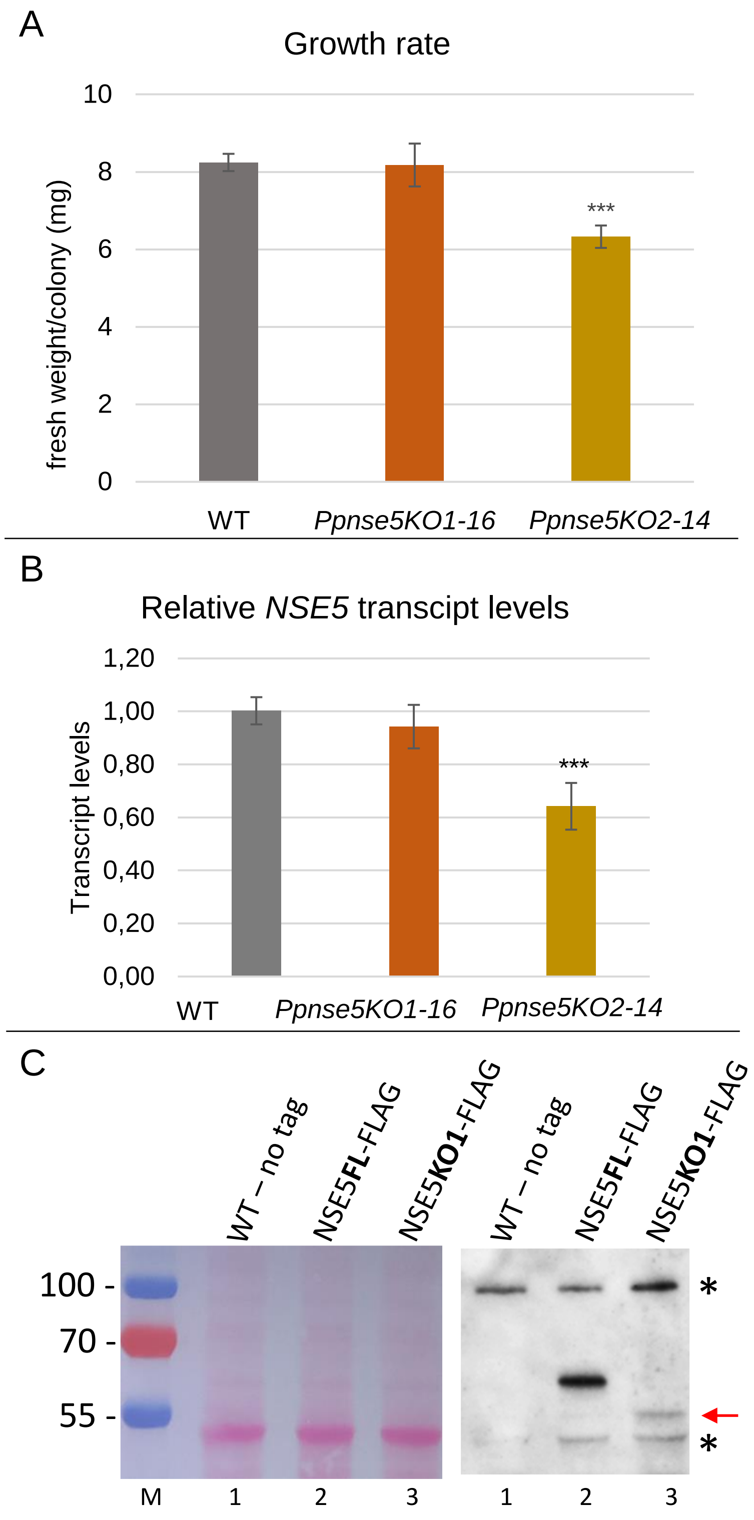
A
Growth rate
fresh weight/colony (mg)
10
8
6
4
2
0
***
WT
Ppnse5KO1-16
Ppnse5KO2-14
B
Relative NSE5 transcipt levels
Transcript levels
1,20
1,00
0,80
0,60
0,40
0,20
0,00
***
WT
Ppnse5KO1-16
Ppnse5KO2-14
C
WT – no tag
NSE5FL-FLAG
NSE5KO1-FLAG
WT – no tag
NSE5FL-FLAG
NSE5KO1-FLAG
100 -
70 -
55 -
*
*
M
1
2
3
1
2
3

**Supporting Table ST1 – List of cloning primers**

| | | | |
|---|---|---|---|
| PpNSE5 (aa1-526) to pGBKT7 | MP441 | fw | 5′ atctcagaggaggacctgcatATGCCTCGCAAAAAGCAG 3′ |
| | MP442 | rev | 5′ ggccgctgcaggtcgacggatccTCAGGAAACTTGCAGGGTA 3′ |
| PpNSE5 (aa190-526) to pGBKT7 | JJ073 | fw | 5′ atctcagaggaggacctgcatatgATGGCTAACGAAAAGCCC 3′ |
| | EB258 | rev | 5′ gcggccgctgcaggtcgacggatccTCAGGAAACTTGCAGGGTATG 3′ |
| PpNSE5 (aa230-526) to pGBKT7 | EB257 | fw | 5′ atctcagaggaggacctgcatATGGCTTCTAAAAATGAAGACCCAC 3′ |
| | EB258 | rev | 5′ gcggccgctgcaggtcgacggatccTCAGGAAACTTGCAGGGTATG 3′ |
| PpNSE5 (aa265-526) to pGBKT7 | JJ084 | fw | 5′ atctcagaggaggacctgcatatgATGCAAGGAAATCCAGAGAATTC 3′ |
| | EB258 | rev | 5′ gcggccgctgcaggtcgacggatccTCAGGAAACTTGCAGGGTATG 3′ |
| PpNSE5 (aa1-505) to pGBKT7 | JJ052 | fw | 5′ atctcagaggaggacctgcatATGATGCCTCGCAAAAAGCAG 3′ |
| | JJ085 | rev | 5′ gcggccgctgcaggtcgacggatccAATCATTTCATCTTCGCATCC 3′ |
| PpNSE5 (aa1-485) to pGBKT7 | JJ052 | fw | 5′ atctcagaggaggacctgcatATGATGCCTCGCAAAAAGCAG 3′ |
| | JJ075 | rev | 5′ gcggccgctgcaggtcgacggatccTCAGTGTGCGTCATGACTTG 3′ |
| PpNSE5 (aa1-455) to pGBKT7 | JJ052 | fw | 5′ atctcagaggaggacctgcatATGATGCCTCGCAAAAAGCAG 3′ |
| | JJ077 | rev | 5′ gcggccgctgcaggtcgacggatccTCATAGGTTAACATTTGAAACGC 3′ |
| PpNSE5 (aa72-526) to pGBKT7 | JJ050 | fw | 5′ atctcagaggaggacctgcatATGATGGGAGCCTCACTAACG 3′ |
| | EB258 | rev | 5′ gcggccgctgcaggtcgacggatccTCAGGAAACTTGCAGGGTATG 3′ |
| PpNSE5 (aa109-526) to pGBKT7 | JJ051 | fw | 5′ atctcagaggaggacctgcatATGATGTCTGTTGTACCAGATG 3′ |
| | EB258 | rev | 5′ gcggccgctgcaggtcgacggatccTCAGGAAACTTGCAGGGTATG 3′ |
| PpNSE5 (aa245-526) to pGBKT7 | EB279 | fw | 5′ atctcagaggaggacctgcatatgATGCTGCGCTGGTTAG 3′ |
| | EB258 | rev | 5′ gcggccgctgcaggtcgacggatccTCAGGAAACTTGCAGGGTATG 3′ |
| PpNSE5 (aa284-526) to pGBKT7 | EB280 | fw | 5′ atctcagaggaggacctgcatatgATGGGGCAAAAAAATGTAAAAATTAAG 3′ |
| | EB258 | rev | 5′ gcggccgctgcaggtcgacggatccTCAGGAAACTTGCAGGGTATG 3′ |
| PpNSE5 (aa1-230) to pGADT7 | EB306 | fw | 5′ gacgtaccagattacgctcatatgATGCCTCGCAAAAAGC 3′ |
| | EB307 | rev | 5′ tctgcagctcgagctcgatggatccTCAAGCAGAGTCAACACCAAAATC 3′ |
| PpNSE5 (aa1-230) to pET-28c(+) | JJ090 | fw | 5′ tggacagcaaatgggtcggatccccATGCCGCGTAAAAAACAG 3´ |
| | JJ089 | rev | 5′ agtggtggtggtggtggtgctcgagTTAAGCAGAGTCAACACC 3´ |
| PpNSE5 (aa72-230) to pET-28c(+) | JJ166 | fw | 5′ ggtggacagcaaatgggtcggatccccATGGGTGCTTCTCTGACC 3´ |
| | JJ089 | rev | 5′ agtggtggtggtggtggtgctcgagTTAAGCAGAGTCAACACC 3´ |
| PpSMC5 (aa280-430) to pGBKT7 | MP414 | fw | 5′ atctcagaggaggacctgcatATGAAGCGTCTGCTTAATGAAG 3′ |
| | BK001 | rev | 5′ tcgacggatccccgggaattccaTGCTATTCTGCGAGTAATTTG 3′ |
| PpSMC5 (aa630-775) to PpSMC5 (aa280-430) | BK002 | fw | 5′ aatagcatggaattcccggggaGACACGAGGAAAAAGAATG 3′ |

| | | | |
|---|---|---|---|
| | EB222 | rev | 5′ gcggccgctgcaggtcgacggatccTTAGTGACTCTTGAGTTCCCTT 3´ |
| PpSMC5 (aa280-380) to pGBKT7 | MP414 | fw | 5′ aatagcatggaattcccggggaGACACGAGGAAAAAGAATG 3′ |
| | BK003 | rev | 5′ gcggccgctgcaggtcgacggatccTTAGTGACTCTTGAGTTCCCTT 3´ |
| PpSMC5 (aa685-775) to PpSMC5 (aa280-380) | BK004 | fw | 5′ aatagcatggaattcccggggaGACACGAGGAAAAAGAATG 3′ |
| | EB222 | rev | 5′ gcggccgctgcaggtcgacggatccTTAGTGACTCTTGAGTTCCCTT 3´ |
| PpSMC6 (aa255-923) to pGBKT7 | BK005 | fw | 5′ aatagcatggaattcccggggaGACACGAGGAAAAAGAATG 3′ |
| | BK006 | rev | 5′ gcggccgctgcaggtcgacggatccTTAGTGACTCTTGAGTTCCCTT 3´ |
| PpSMC6 (aa226-405) to pGBKT7 | EB259 | fw | 5′ aatagcatggaattcccggggaGACACGAGGAAAAAGAATG 3′ |
| | BK007 | rev | 5′ gcggccgctgcaggtcgacggatccTTAGTGACTCTTGAGTTCCCTT 3´ |
| PpSMC6 (aa750-955) to PpSMC6 (aa226-405) | BK008 | fw | 5′ aatagcatggaattcccggggaGACACGAGGAAAAAGAATG 3′ |
| | EB260 | rev | 5′ gcggccgctgcaggtcgacggatccTTAGTGACTCTTGAGTTCCCTT 3´ |
| PpSMC6 (aa226-310) to pGBKT7 | EB259 | fw | 5′ aatagcatggaattcccggggaGACACGAGGAAAAAGAATG 3′ |
| | EB242 | rev | 5′ gcggccgctgcaggtcgacggatccTTAGTGACTCTTGAGTTCCCTT 3´ |
| PpSMC6 (aa836-955) to PpSMC6 (aa226-310) | EB243 | fw | 5′ aatagcatggaattcccggggaGACACGAGGAAAAAGAATG 3′ |
| | EB260 | rev | 5′ gcggccgctgcaggtcgacggatccTTAGTGACTCTTGAGTTCCCTT 3´ |

### Supporting Table ST2: Primers for moss analysis and manipulation

| | | | |
|---|---|---|---|
| donor template PpNSE5KO1 | pKA1390 | fw | 5’ CCCGACGATTAGTCATACTGAGACCGCTGCTCAACCTGCCACCCG 3’ |
| | pKA1391 | rev | 5’ CGGGTGGCAGGTTGAGCAGCGGTCTCAGTATGACTAATCGTCGGG 3’ |
| sgRNA PpNSE5KO1 | pKA1392 | fw | 5’ CCATGCAGCAGTCTCAGTAGATGA 3’ |
| | pKA1393 | rev | 5’ AAACTCATCTACTGAGACTGCTGC 3’ |
| donor template PpNSE5KO2 | pKA1394 | fw | 5’ GGGAAAAGCTAAGTGTACACTTGTCGACAATAAGTAATA 3’ |
| | pKA1395 | rev | 5’ TATTACTTATTGTCGACAAGTGTACACTTAGCTTTTCCCA 3’ |
| sgRNA PpNSE5KO2 | pKA1396 | fw | 5’ CCATGTACGAACTTGTAGACAATA 3’ |
| | pKA1397 | rev | 5’ AAACTATTGTCTACAAGTTCGTAC 3’ |
| PpNSE5KO1 verification | pKA1437 | fw | 5’ GCACCCTTCAGATCTTTCTC 3’ |
| | pKA1438 | rev | 5’ GTGCAGAGCTTCTGGTGATG 3’ |
| PpNSE5KO2 verification | pKA1439 | fw | 5’ TGACCTCGATCTGGTTAGTG 3’ |
| | pKA1440 | rev | 5’ CCCGAGAGTTAACGTACAAG 3’ |
| PpNSE5, transcript | pKA1292 | fw | 5’ TTATGGGAGCCTCACTAACG 3’ |
| | pKA1293 | rev | 5’ AACAGACATCACCCGAACAG 3’ |
| sgRNA NSE5-FLAG | pKA1500 | fw | 5’ GTGGAGGACTTACAACCCATGCAGGGTATGAGCCATGACCGTTTTAGAGCTATGCTGAAA 3’ |
| | pKA1501 | rev | 5’ TTTCAGCATAGCTCTAAAACGGTCATGGCTCATACCCTGCATGGGTTGTAAGTCCTCCAC 3’ |
| NSE5 3'end targeting arm | pKA1493 | fw | 5’TATCGATAAGCTTGATATCGGAAGTTACACATCATACTGCGAGATGTGCACTGGTCATGGTATCGATAAGCTTGATATCGG<br>AAGTTACACATCATACTGCGAGATGTGCACTGGTCATGG 3’ |
| | pKA1494 | rev | 5’TGGATCCCCCGGGCTGCAGGGGAAACTTGCAGGGTATGTGCCATGACCAGTGCACATCTCGCAGTATGATGTGTAACTTC<br>CGATATCAAGCTTATCGATA 3’ |
| NSE5 3'UTR targeting arm | pKA1498 | fw | 5’TGACAAGTAGGAAGGAAATTTGCTTGTACCATTTACGCATATCTAGGTACAAAACGATCACTGCATTTTCCCTGCAGCCCG<br>GGGGATCCA 3’ |
| | pKA1499 | rev | 5’TGGATCCCCCGGGCTGCAGGGAAAATGCAGTGATCGTTTTGTACCTAGATATGCGTAAATGGTACAAGCAAATTTCCTTCC<br>TACTTGTCA 3’ |
| FLAG-tag amplification | pKA1496 | fw | 5’ GCAAGTTTCCCATATGTCGTACGCTGCAG3’ |
| | pKA1497 | rev | 5’ AATTTCCTTCCTACTTGTCATCGCCATC 3’ |
| FLAG construct amplification | pKA1502 | fw | 5’ GAAGTTACACATCATACTGC 3’ |
| | pKA1503 | rev | 5’ AAAATGCAGTGATCGTTTTG 3’ |
| NSE5-FLAG verification | pKA1408 | fw | 5’GAAACAGGCTAAGGGTCTTC 3’ |
| | pKA1470 | rev | 5’ GTCAGTTCTTTGCCAGTCAC 3’ |
